## Supplemental figures for "Antibody evasion by SARS-CoV-2 Omicron subvariants BA.2.12.1, BA.4, and BA.5"

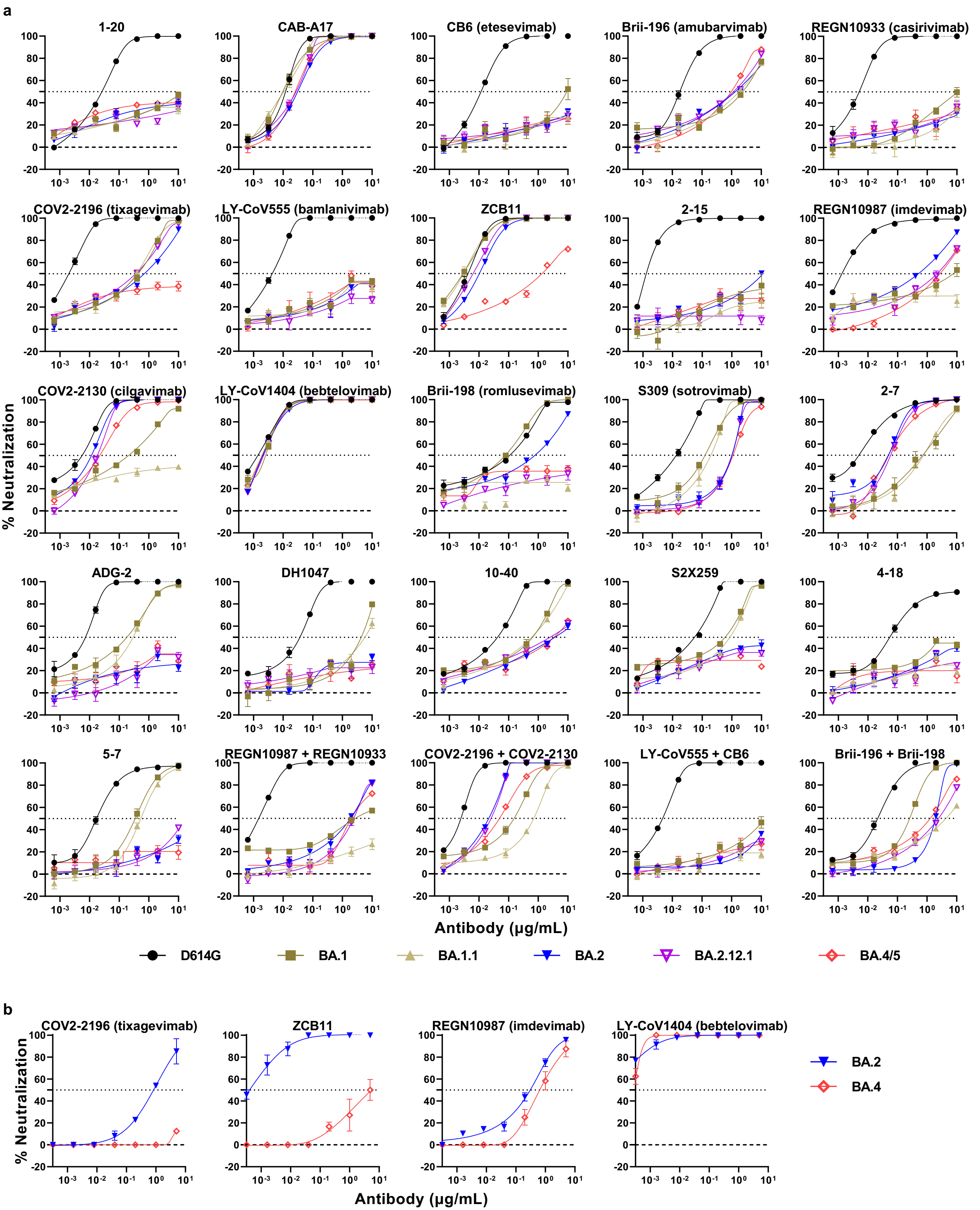
a
1-20
CAB-A17
CB6 (etesevimab)
Brii-196 (amubarvimab)
REGN10933 (casirivimab)
COV2-2196 (tixagevimab)
LY-CoV555 (bamlanivimab)
ZCB11
2-15
REGN10987 (imdevimab)
COV2-2130 (cilgavimab)
LY-CoV1404 (bebtelovimab)
Brii-198 (romlusevimab)
S309 (sotrovimab)
2-7
ADG-2
DH1047
10-40
S2X259
4-18
5-7
REGN10987 + REGN10933
COV2-2196 + COV2-2130
LY-CoV555 + CB6
Brii-196 + Brii-198
% Neutralization
Antibody (μg/mL)
D614G
BA.1
BA.1.1
BA.2
BA.2.12.1
BA.4/5
b
COV2-2196 (tixagevimab)
ZCB11
REGN10987 (imdevimab)
LY-CoV1404 (bebtelovimab)
% Neutralization
Antibody (μg/mL)
BA.2
BA.4

**a**

| $IC_{50}$ (µg/mL) | RBD mAbs | | | | | | | | | | | | | | | | | | | | NTD mAbs | | Combination | | |
|---|---|---|---|---|---|---|---|---|---|---|---|---|---|---|---|---|---|---|---|---|---|---|---|---|---|
| | Class 1 | | | | Class 2 | | | | | Class 3 | | | | | | Class 4 | | | | | | | | | |
| | 1-20 | CAB-A17 | CB6 | Brii-196 | REGN 10933 | COV2-2196 | LY-CoV555 | ZCB11 | 2-15 | REGN 10987 | COV2-2130 | LY-CoV 1404 | Brii-198 | S309 | 2-7 | ADG-2 | DH1047 | 10-40 | S2X259 | 4-18 | 5-7 | REGN 10987 + REGN 10933 | COV2-2196 + COV2-2130 | LY-CoV555 + CB6 | Brii-196 + Brii-198 |
| D614G | 0.027 | 0.012 | 0.012 | 0.018 | 0.005 | 0.002 | 0.004 | 0.004 | 0.001 | 0.001 | 0.006 | 0.001 | 0.110 | 0.014 | 0.005 | 0.007 | 0.037 | 0.042 | 0.055 | 0.054 | 0.017 | 0.001 | 0.002 | 0.005 | 0.022 |
| BA.1 | >10 | 0.010 | 9.253 | 2.385 | >10 | 0.432 | >10 | 0.003 | >10 | 7.586 | 0.209 | 0.003 | 0.078 | 0.127 | 0.716 | 0.181 | 4.322 | 0.644 | 0.590 | >10 | 0.347 | 2.951 | 0.154 | >10 | 0.260 |
| BA.1.1 | >10 | 0.010 | >10 | 1.792 | >10 | 0.385 | >10 | 0.003 | >10 | >10 | >10 | 0.002 | >10 | 0.200 | 0.763 | 0.295 | 5.723 | 0.600 | 0.859 | >10 | 0.534 | >10 | 0.708 | >10 | 4.394 |
| BA.2 | >10 | 0.031 | >10 | 1.346 | >10 | 0.704 | >10 | 0.010 | >10 | 0.505 | 0.012 | 0.002 | 0.782 | 1.019 | 0.050 | >10 | >10 | 3.642 | >10 | >10 | >10 | 1.882 | 0.021 | >10 | 1.907 |
| BA.2.12.1 | >10 | 0.027 | >10 | 1.171 | >10 | 0.361 | >10 | 0.006 | >10 | 2.125 | 0.018 | 0.002 | >10 | 1.035 | 0.059 | >10 | >10 | 2.519 | >10 | >10 | >10 | 2.400 | 0.026 | >10 | 1.936 |
| BA.4/5 | >10 | 0.025 | >10 | 0.978 | >10 | >10 | >10 | 1.351 | >10 | 2.682 | 0.022 | 0.002 | >10 | 1.120 | 0.049 | >10 | >10 | 3.404 | >10 | >10 | >10 | 1.998 | 0.049 | >10 | 2.445 |

<0.01 <0.1 <1 <10 >10

**b**

| $IC_{50}$ (µg/mL) | RBD mAbs | | | |
|---|---|---|---|---|
| | Class 2 | | Class 3 | |
| | COV2-2196 | ZCB11 | REGN 10987 | LY-CoV1404 |
| BA.2 | 0.8287 | 0.0004 | 0.3057 | <0.001 |
| BA.4 | >5 | >5 | 0.6418 | <0.001 |

<0.01 <0.1 <1 <5 >5

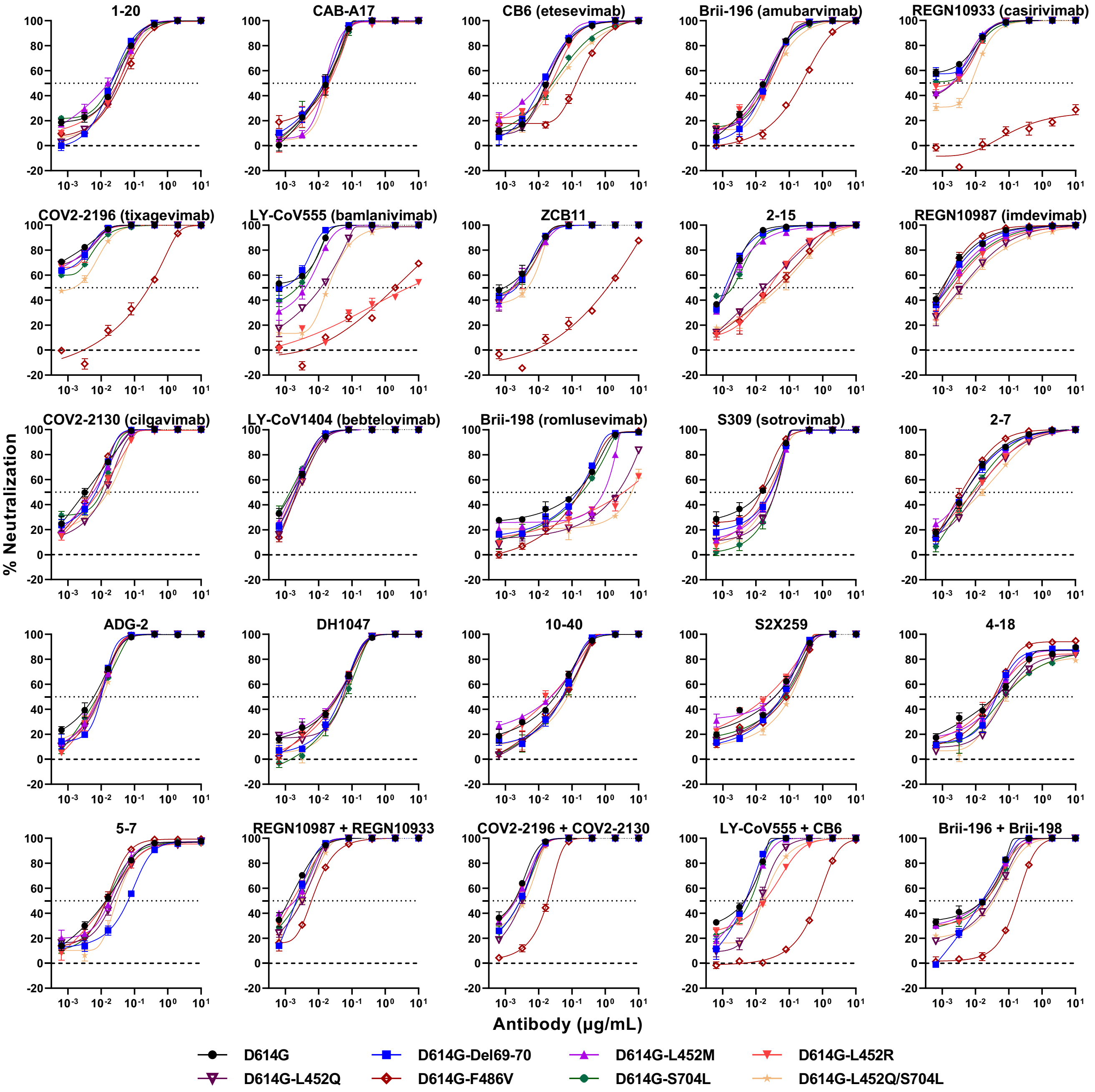

1-20
CAB-A17
CB6 (etesevimab)
Brii-196 (amubarvimab)
REGN10933 (casirivimab)
COV2-2196 (tixagevimab)
LY-CoV555 (bamlanivimab)
ZCB11
2-15
REGN10987 (imdevimab)
COV2-2130 (cilgavimab)
LY-CoV1404 (bebtelovimab)
Brii-198 (romlusevimab)
S309 (sotrovimab)
2-7
ADG-2
DH1047
10-40
S2X259
4-18
5-7
REGN10987 + REGN10933
COV2-2196 + COV2-2130
LY-CoV555 + CB6
Brii-196 + Brii-198
% Neutralization
Antibody (μg/mL)
D614G
D614G-Del69-70
D614G-L452M
D614G-L452R
D614G-L452Q
D614G-F486V
D614G-S704L
D614G-L452Q/S704L

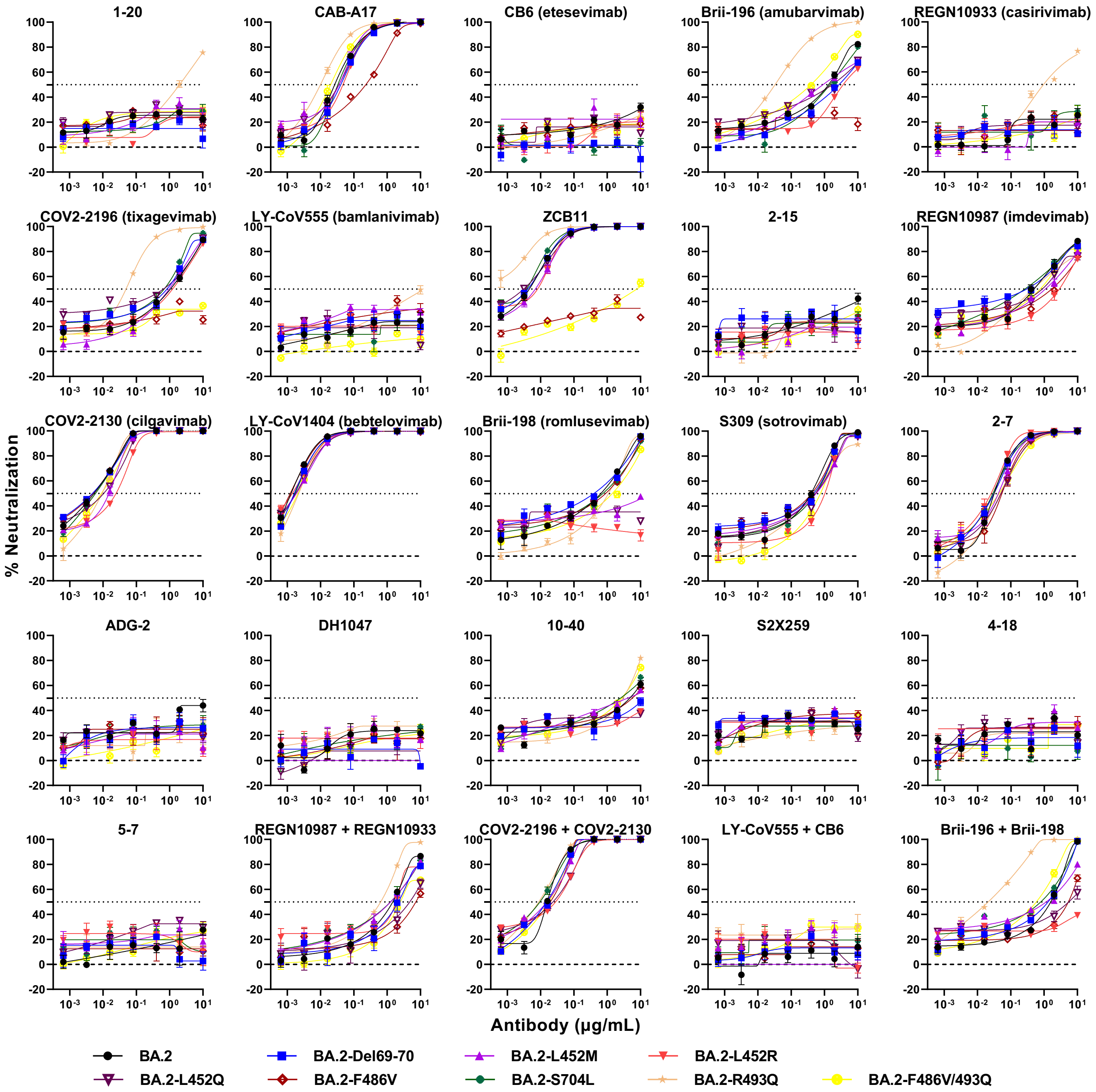

1-20
CAB-A17
CB6 (etesevimab)
Brii-196 (amubarvimab)
REGN10933 (casirivimab)
COV2-2196 (tixagevimab)
LY-CoV555 (bamlanivimab)
ZCB11
2-15
REGN10987 (imdevimab)
COV2-2130 (cilgavimab)
LY-CoV1404 (bebtelovimab)
Brii-198 (romlusevimab)
S309 (sotrovimab)
2-7
ADG-2
DH1047
10-40
S2X259
4-18
5-7
REGN10987 + REGN10933
COV2-2196 + COV2-2130
LY-CoV555 + CB6
Brii-196 + Brii-198
% Neutralization
Antibody (μg/mL)
BA.2
BA.2-Del69-70
BA.2-L452M
BA.2-L452R
BA.2-L452Q
BA.2-F486V
BA.2-S704L
BA.2-R493Q
BA.2-F486V/493Q

**a**

| $IC_{50}$ (μg/mL) | RBD mAbs | | | | | | | | | | | | | | | | | | | NTD mAbs | | Combination | | | |
|---|---|---|---|---|---|---|---|---|---|---|---|---|---|---|---|---|---|---|---|---|---|---|---|---|---|
| | Class 1 | | | | Class 2 | | | | | Class 3 | | | | | | Class 4 | | | | | | | | | |
| | 1-20 | CAB-A17 | CB6 | Brii-196 | REGN 10933 | COV2-2196 | LY-CoV 555 | ZCB11 | 2-15 | REGN 10987 | COV2-2130 | LY-CoV 1404 | Brii-198 | S309 | 2-7 | ADG-2 | DH1047 | 10-40 | S2X259 | 4-18 | 5-7 | REGN 10987 + REGN 10933 | COV2-2196 + COV2-2130 | LY-CoV555 + CB6 | Brii-196 + Brii-198 |
| D614G | 0.026 | 0.016 | 0.017 | 0.016 | <0.001 | <0.001 | 0.002 | 0.002 | 0.002 | 0.001 | 0.003 | 0.002 | 0.129 | 0.014 | 0.005 | 0.006 | 0.037 | 0.031 | 0.036 | 0.038 | 0.014 | 0.001 | 0.002 | 0.005 | 0.016 |
| D614G-Del69-70 | 0.020 | 0.014 | 0.013 | 0.021 | <0.001 | <0.001 | 0.001 | 0.002 | 0.001 | 0.001 | 0.007 | 0.002 | 0.148 | 0.028 | 0.005 | 0.010 | 0.047 | 0.051 | 0.059 | 0.042 | 0.064 | 0.002 | 0.003 | 0.005 | 0.016 |
| D614G-L452M | 0.017 | 0.016 | 0.011 | 0.016 | 0.002 | <0.001 | 0.004 | 0.002 | 0.001 | 0.001 | 0.006 | 0.002 | 0.892 | 0.029 | 0.005 | 0.009 | 0.034 | 0.023 | 0.036 | 0.042 | 0.018 | 0.002 | 0.002 | 0.005 | 0.016 |
| D614G-L452R | 0.032 | 0.020 | 0.024 | 0.024 | 0.002 | <0.001 | 5.018 | 0.002 | 0.024 | 0.002 | 0.009 | 0.002 | 3.526 | 0.023 | 0.009 | 0.008 | 0.039 | 0.023 | 0.020 | 0.035 | 0.014 | 0.002 | 0.002 | 0.018 | 0.021 |
| D614G-L452Q | 0.033 | 0.014 | 0.018 | 0.023 | 0.002 | <0.001 | 0.012 | 0.002 | 0.017 | 0.004 | 0.013 | 0.002 | 2.346 | 0.038 | 0.011 | 0.009 | 0.056 | 0.055 | 0.061 | 0.078 | 0.022 | 0.003 | 0.003 | 0.013 | 0.031 |
| D614G-F486V | 0.039 | 0.019 | 0.135 | 0.231 | >10 | 0.272 | 1.961 | 1.174 | 0.036 | 0.001 | 0.005 | 0.002 | 0.175 | 0.014 | 0.004 | 0.009 | 0.033 | 0.051 | 0.079 | 0.034 | 0.014 | 0.006 | 0.019 | 0.701 | 0.174 |
| D614G-S704L | 0.020 | 0.017 | 0.026 | 0.019 | <0.001 | <0.001 | 0.002 | 0.002 | 0.002 | 0.002 | 0.010 | 0.001 | 0.199 | 0.038 | 0.008 | 0.008 | 0.061 | 0.057 | 0.069 | 0.071 | 0.015 | 0.002 | 0.003 | 0.007 | 0.019 |
| D614G-L452Q/S704L | 0.033 | 0.020 | 0.032 | 0.026 | 0.007 | 0.001 | 0.019 | 0.005 | 0.049 | 0.004 | 0.016 | 0.002 | 6.166 | 0.028 | 0.015 | 0.011 | 0.052 | 0.074 | 0.106 | 0.076 | 0.029 | 0.003 | 0.004 | 0.016 | 0.035 |

**b**

| $IC_{50}$ (μg/mL) | RBD mAbs | | | | | | | | | | | | | | | | | | | NTD mAbs | | Combination | | | |
|---|---|---|---|---|---|---|---|---|---|---|---|---|---|---|---|---|---|---|---|---|---|---|---|---|---|
| | Class 1 | | | | Class 2 | | | | | Class 3 | | | | | | Class 4 | | | | | | | | | |
| | 1-20 | CAB-A17 | CB6 | Brii-196 | REGN 10933 | COV2-2196 | LY-CoV 555 | ZCB11 | 2-15 | REGN 10987 | COV2-2130 | LY-CoV 1404 | Brii-198 | S309 | 2-7 | ADG-2 | DH1047 | 10-40 | S2X259 | 4-18 | 5-7 | REGN 10987 + REGN 10933 | COV2-2196 + COV2-2130 | LY-CoV55 5 + CB6 | Brii-196 + Brii-198 |
| BA.2 | >10 | 0.027 | >10 | 1.329 | >10 | 1.060 | >10 | 0.005 | >10 | 0.495 | 0.005 | 0.001 | 0.642 | 0.393 | 0.032 | >10 | >10 | 4.824 | >10 | >10 | >10 | 1.475 | 0.016 | >10 | 1.592 |
| BA.2-Del69-70 | >10 | 0.040 | >10 | 2.726 | >10 | 0.835 | >10 | 0.004 | >10 | 0.298 | 0.005 | 0.002 | 0.394 | 0.469 | 0.031 | >10 | >10 | >10 | >10 | >10 | >10 | 2.178 | 0.015 | >10 | 1.320 |
| BA.2-L452M | >10 | 0.036 | >10 | 0.907 | >10 | 0.970 | >10 | 0.007 | >10 | 1.081 | 0.015 | 0.002 | >10 | 0.557 | 0.042 | >10 | >10 | 4.246 | >10 | >10 | >10 | 1.276 | 0.015 | >10 | 1.163 |
| BA.2-L452R | >10 | 0.047 | >10 | 6.815 | >10 | 1.228 | >10 | 0.008 | >10 | 2.832 | 0.025 | 0.001 | >10 | 1.022 | 0.026 | >10 | >10 | >10 | >10 | >10 | >10 | 1.864 | 0.028 | >10 | >10 |
| BA.2-L452Q | >10 | 0.036 | >10 | 1.717 | >10 | 0.655 | >10 | 0.003 | >10 | 0.872 | 0.010 | 0.002 | >10 | 0.535 | 0.051 | >10 | >10 | >10 | >10 | >10 | >10 | 4.793 | 0.024 | >10 | 5.525 |
| BA.2-F486V | >10 | 0.229 | >10 | >10 | >10 | >10 | >10 | >10 | >10 | 1.681 | 0.005 | 0.001 | 0.887 | 0.412 | 0.054 | >10 | >10 | 5.759 | >10 | >10 | >10 | 7.366 | 0.020 | >10 | 5.377 |
| BA.2-R493Q | 2.020 | 0.010 | >10 | 0.033 | 0.960 | 0.049 | >10 | <0.001 | >10 | 0.454 | 0.009 | 0.002 | 1.089 | 0.485 | 0.049 | >10 | >10 | 3.008 | >10 | >10 | >10 | 0.641 | 0.009 | >10 | 0.021 |
| BA.2-S704L | >10 | 0.033 | >10 | 1.464 | >10 | 0.686 | >10 | 0.004 | >10 | 0.262 | 0.006 | 0.002 | 0.735 | 0.539 | 0.029 | >10 | >10 | 2.537 | >10 | >10 | >10 | 1.262 | 0.010 | >10 | 0.800 |
| BA.2-F486V/R493Q | >10 | 0.020 | >10 | 0.394 | >10 | >10 | >10 | 7.766 | >10 | 0.757 | 0.009 | 0.002 | 1.414 | 0.754 | 0.044 | >10 | >10 | 2.751 | >10 | >10 | >10 | 2.498 | 0.017 | >10 | 0.586 |

| <0.01 | <0.1 | <1 | <10 | >10 |
|---|---|---|---|---|

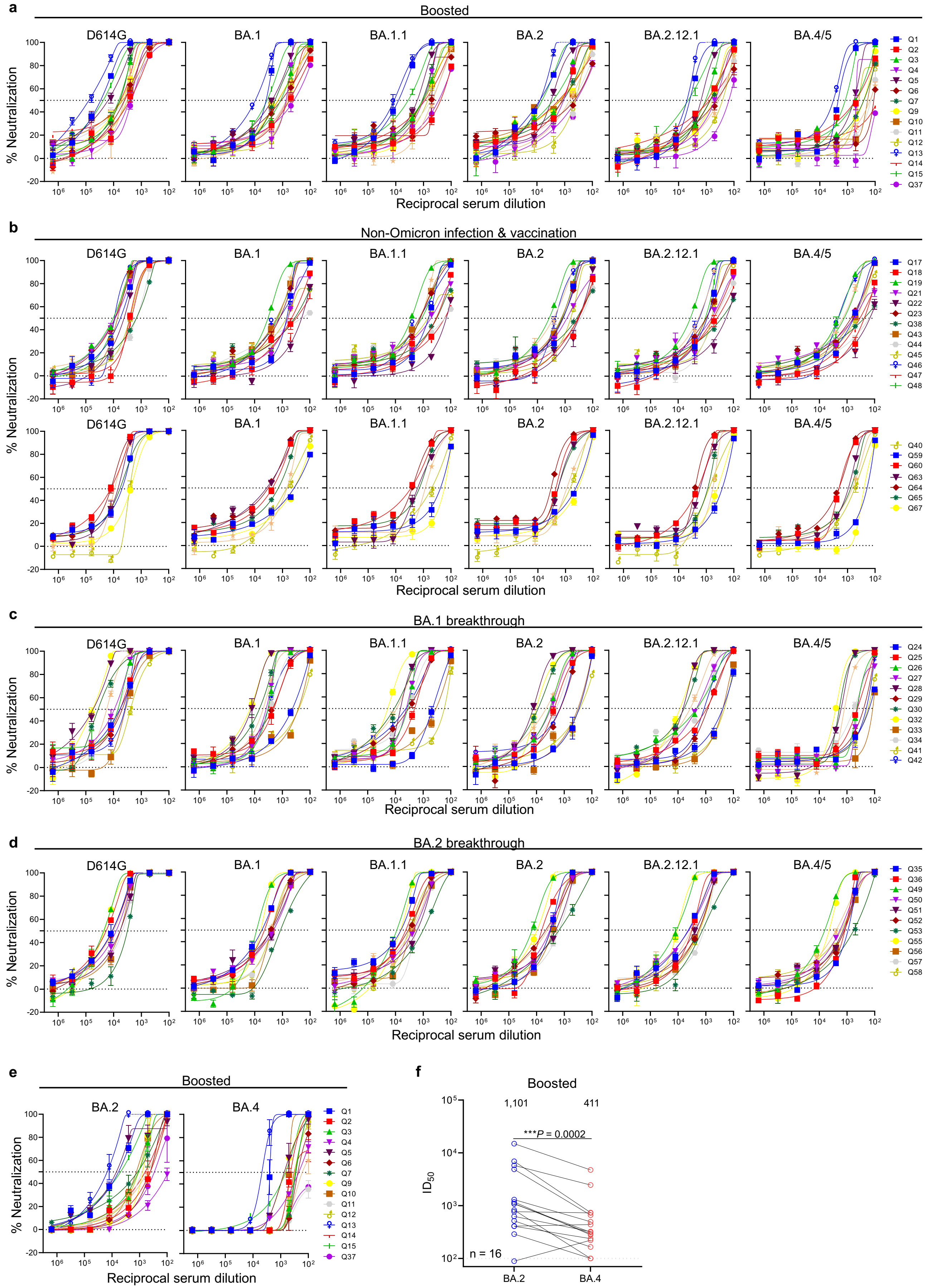
a
Boosted
D614G
BA.1
BA.1.1
BA.2
BA.2.12.1
BA.4/5
% Neutralization
Reciprocal serum dilution
Q1 Q2 Q3 Q4 Q5 Q6 Q7 Q9 Q10 Q11 Q12 Q13 Q14 Q15 Q37
b
Non-Omicron infection & vaccination
Q17 Q18 Q19 Q21 Q22 Q23 Q38 Q43 Q44 Q45 Q46 Q47 Q48
Q40 Q59 Q60 Q63 Q64 Q65 Q67
c
BA.1 breakthrough
Q24 Q25 Q26 Q27 Q28 Q29 Q30 Q32 Q33 Q34 Q41 Q42
d
BA.2 breakthrough
Q35 Q36 Q49 Q50 Q51 Q52 Q53 Q55 Q56 Q57 Q58
e
Boosted
BA.2
BA.4
f
Boosted
1,101
411
***P = 0.0002
ID50
n = 16
BA.2
BA.4

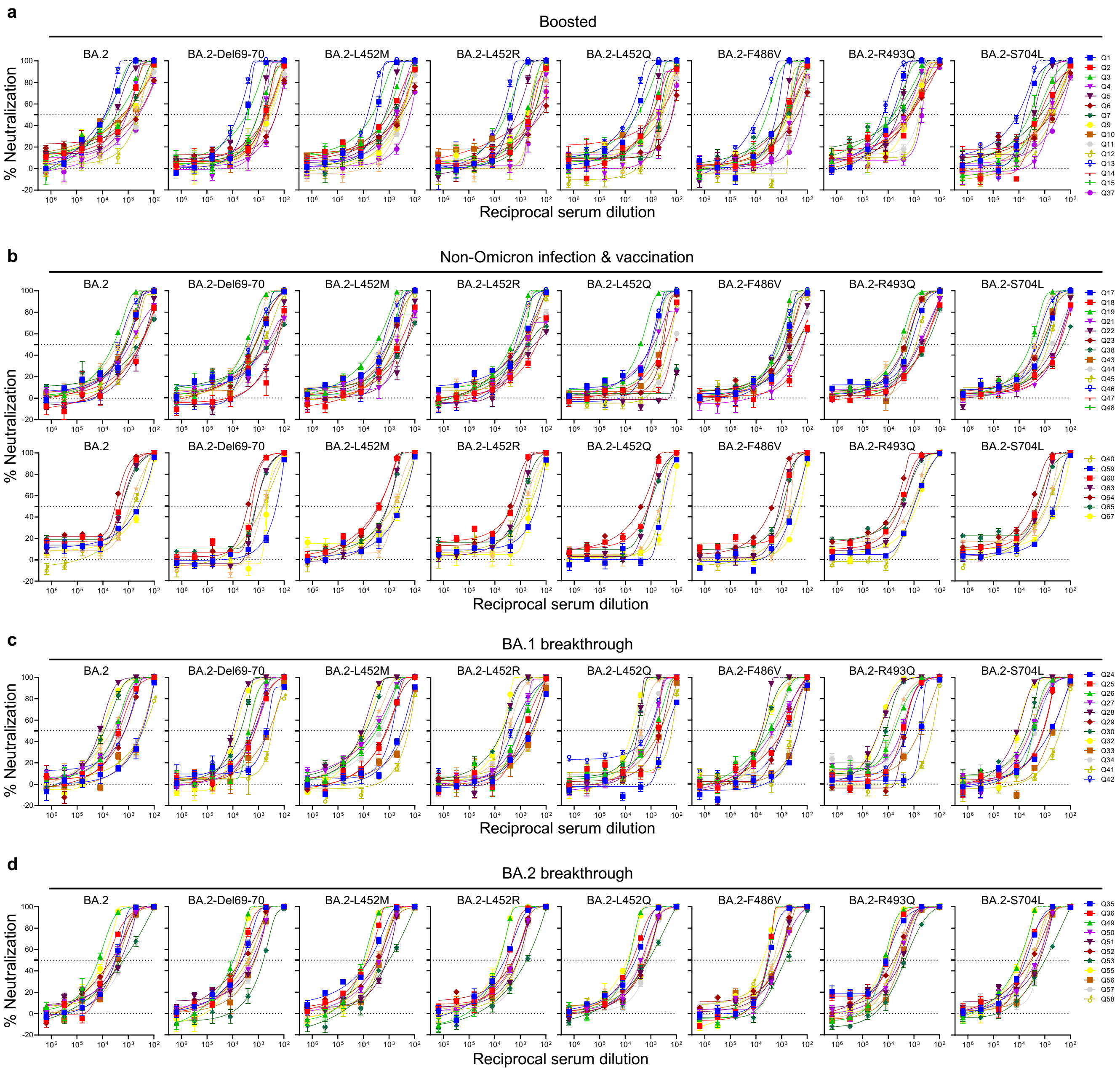
a
Boosted
BA.2
BA.2-Del69-70
BA.2-L452M
BA.2-L452R
BA.2-L452Q
BA.2-F486V
BA.2-R493Q
BA.2-S704L
% Neutralization
Reciprocal serum dilution
Q1 Q2 Q3 Q4 Q5 Q6 Q7 Q9 Q10 Q11 Q12 Q13 Q14 Q15 Q37
b
Non-Omicron infection & vaccination
Q17 Q18 Q19 Q21 Q22 Q23 Q38 Q43 Q44 Q45 Q46 Q47 Q48
Q40 Q59 Q60 Q63 Q64 Q65 Q67
c
BA.1 breakthrough
Q24 Q25 Q26 Q27 Q28 Q29 Q30 Q32 Q33 Q34 Q41 Q42
d
BA.2 breakthrough
Q35 Q36 Q49 Q50 Q51 Q52 Q53 Q55 Q56 Q57 Q58

| Mutation | Count in BA.1 | Frequency in BA.1 | Count in BA.2 | Frequency in BA.2 | Count in other variants | Frequency in other variants |
|---|---|---|---|---|---|---|
| F486V | 23 | 2.17E-06 | 134 | 1.26E-05 | 898 | 8.48E-05 |
| Del486 | 193 | 1.82E-05 | 549 | 5.18E-05 | 760 | 7.17E-05 |
| F486L | 37 | 3.49E-06 | 10 | 9.44E-07 | 155 | 1.46E-05 |
| F486S | 61 | 5.76E-06 | 10 | 9.44E-07 | 142 | 1.34E-05 |
| F486I | 5 | 4.72E-07 | 2 | 1.89E-07 | 34 | 3.21E-06 |
| F486Y | 12 | 1.13E-06 | 2 | 1.89E-07 | 20 | 1.89E-06 |
| F486W | 8 | 7.55E-07 | 1 | 9.44E-08 | 10 | 9.44E-07 |
| F486T | 5 | 4.72E-07 | 0 | 0 | 5 | 4.72E-07 |
| F486E | 2 | 1.89E-07 | 0 | 0 | 3 | 2.83E-07 |
| F486N | 2 | 1.89E-07 | 0 | 0 | 3 | 2.83E-07 |
| F486H | 2 | 1.89E-07 | 0 | 0 | 2 | 1.89E-07 |
| F486P | 2 | 1.89E-07 | 0 | 0 | 2 | 1.89E-07 |
| F486R | 1 | 9.44E-08 | 0 | 0 | 2 | 1.89E-07 |
| F486C | 0 | 0 | 0 | 0 | 1 | 9.44E-08 |
| F486G | 1 | 9.44E-08 | 0 | 0 | 1 | 9.44E-08 |
| F486M | 0 | 0 | 0 | 0 | 1 | 9.44E-08 |
| F486Q | 0 | 0 | 1 | 9.44E-08 | 1 | 9.44E-08 |

| Sample ID | Vaccine type and infected strain | Days post-vaccination or *infection (after last exposure) | Documented COVID-19 | Age | Gender |
|---|---|---|---|---|---|
| Boosted | | | | | |
| Q1 | mRNA-1273/mRNA-1273/mRNA-1273 | 29 | No | 66 | Female |
| Q2 | BNT162b2/BNT162b2/BNT162b2 | 30 | No | 68 | Male |
| Q3 | BNT162b2/BNT162b2/BNT162b2 | 14 | No | 64 | Female |
| Q4 | BNT162b2/BNT162b2/BNT162b2 | 34 | No | 55 | Male |
| Q5 | BNT162b2/BNT162b2/BNT162b2 | 34 | No | 45 | Male |
| Q6 | BNT162b2/BNT162b2/BNT162b2 | 15 | No | 50 | Female |
| Q7 | BNT162b2/BNT162b2/BNT162b2 | 15 | No | 48 | Female |
| Q8 | BNT162b2/BNT162b2/BNT162b2 | 29 | No | 71 | Male |
| Q9 | BNT162b2/BNT162b2/BNT162b2 | 90 | No | 59 | Male |
| Q10 | BNT162b2/BNT162b2/BNT162b2 | 33 | No | 45 | Male |
| Q11 | BNT162b2/BNT162b2/BNT162b2 | 87 | No | 66 | Female |
| Q12 | BNT162b2/BNT162b2/BNT162b2 | 84 | No | 26 | Male |
| Q13 | mRNA-1273/mRNA-1273/mRNA-1273 | 23 | No | 28 | Female |
| Q14 | BNT162b2/BNT162b2/BNT162b2 | 14 | No | 78 | Male |
| Q15 | BNT162b2/BNT162b2/mRNA-1273 | 32 | No | 39 | Male |
| Q37 | BNT162b2/BNT162b2/BNT162b2 | 20 | No | Unknown | Female |
| Non-Omicron infection & vaccination | | | | | |
| Q17 | R.1/mRNA-1273/mRNA-1273 | 7 | Yes | 34 | Female |
| Q18 | R.1/mRNA-1273/mRNA-1273 | 28 | Yes | 52 | Male |
| Q19 | R.1/mRNA-1273/mRNA-1273 | 21 | Yes | 67 | Female |
| Q21 | R.1/mRNA-1273/mRNA-1273 | >28 | Yes | 57 | Female |
| Q22 | BNT162b2/B.1.526 | *89 | Yes | 42 | Male |
| Q23 | BNT162b2/B.1.526 | *82 | Yes | 32 | Male |
| Q38 | BNT162b2/B.1.1.7 | *59 | Yes | 22 | Female |
| Q39 | BNT162b2/B.1.1.7 | *213 | Yes | 66 | Male |
| Q40 | BNT162b2/B.1.617.2 | *31 | Yes | 50 | Female |
| Q43 | BNT162b2/BNT162b2/B.1.526 | *62 | Yes | 30 | Male |
| Q44 | WA1/mRNA-1273/mRNA-1273 | 114 | Yes | 49 | Female |
| Q45 | WA1/BNT162b2/BNT162b2 | 57 | Yes | 35 | Female |
| Q46 | WA1/BNT162b2/BNT162b2 | 46 | Yes | 30 | Female |
| Q47 | WA1/BNT162b2/BNT162b2 | 57 | Yes | 32 | Female |
| Q48 | WA1/BNT162b2/BNT162b2 | 50 | Yes | 64 | Female |
| Q59 | BNT162b2/BNT162b2/B.1.617.2 | *35 | Yes | 58 | Female |
| Q60 | B.1.617.2/BNT162b2/BNT162b2 | 40 | Yes | 61 | Male |
| Q63 | BNT162b2/BNT162b2/B.1.617.2 | *30 | Yes | 40 | Female |
| Q64 | mRNA-1273/mRNA-1273/B.1.617.2 | *66 | Yes | 29 | Male |
| Q65 | BNT162b2/BNT162b2/B.1.617.2 | *62 | Yes | 33 | Female |
| Q66 | BNT162b2/BNT162b2/B.1.617.2 | *60 | Yes | 42 | Female |
| Q67 | BNT162b2/BNT162b2/B.1.617.2 | *73 | Yes | 37 | Male |
| BA.1 breakthrough | | | | | |
| Q24 | BNT162b2/BNT162b2/BA.1 | *14 | Yes | Unknown | Unknown |
| Q25 | BNT162b2/BNT162b2/BA.1 | *14 | Yes | Unknown | Unknown |
| Q26 | mRNA-1273/mRNA-1273/BA.1 | *35 | Yes | Unknown | Unknown |
| Q27 | BNT162b2/BNT162b2/BNT162b2/BA.1 | *135 | Yes | 78 | Male |
| Q28 | BNT162b2/BNT162b2/BNT162b2/BA.1 | *14 | Yes | Unknown | Unknown |
| Q29 | BNT162b2/BNT162b2/BNT162b2/BA.1 | *14 | Yes | Unknown | Unknown |
| Q30 | BNT162b2/BNT162b2/BNT162b2/BA.1 | *14 | Yes | Unknown | Unknown |
| Q31 | BNT162b2/BNT162b2/BNT162b2/BA.1 | *41 | Yes | 48 | Male |
| Q32 | BNT162b2/BNT162b2/BNT162b2/BA.1 | *26 | Yes | 38 | Female |
| Q33 | BNT162b2/BNT162b2/B.1.617.2/BNT162b2/BA.1 | *19 | Yes | 35 | Female |
| Q34 | BNT162b2/BNT162b2/mRNA-1273/mRNA-1273/BA.1 | *67 | Yes | 40 | Male |
| Q41 | WA1/BNT162b2/BA.1 | *21 | Yes | 52 | Male |
| Q42 | WA1/BNT162b2/BA.1 | *44 | Yes | 37 | Intersex |
| BA.2 breakthrough | | | | | |
| Q35 | BNT162b2/BNT162b2/BA.2 | *14 | Yes | 50 | Female |
| Q36 | BNT162b2/BNT162b2/BNT162b2/Ad26.COV2.S/BA.2 | *22 | Yes | 69 | Male |
| Q49 | BNT162b2/BNT162b2/mRNA-1273/BA.2 | *16 | Yes | 32 | Male |
| Q50 | mRNA-1273/mRNA-1273/mRNA-1273/BA.2 | *14 | Yes | 34 | Male |
| Q51 | BNT162b2/BNT162b2/mRNA-1273/BA.2 | *19 | Yes | 33 | Female |
| Q52 | BNT162b2/BNT162b2/mRNA-1273/BA.2 | *18 | Yes | 29 | Female |
| Q53 | BNT162b2/BNT162b2/BNT162b2/BA.2 | *25 | Yes | 34 | Male |
| Q54 | BNT162b2/BNT162b2/BNT162b2/BA.2 | *36 | Yes | 37 | Female |
| Q55 | BNT162b2/BNT162b2/mRNA-1273/BA.2 | *18 | Yes | 41 | Female |
| Q56 | mRNA-1273/mRNA-1273/mRNA-1273/BA.2 | *21 | Yes | 36 | Female |
| Q57 | BNT162b2/BNT162b2/mRNA-1273/BA.2 | *32 | Yes | 28 | Male |
| Q58 | BNT162b2/BNT162b2/mRNA-1273/BA.2 | *23 | Yes | 33 | Female |
